# GeoEPred: A Multimodal Structure-Aware Geometric Deep Learning Framework for Gram-Negative Bacterial Secreted Effector Prediction with Sequence Semantics

**DOI:** 10.64898/2026.05.18.725929

**Authors:** Shouzhen Song, Hua Shi, Hongfeng Wu, Dachen Liu, Yihang Lin, Nor Ashidi Mat Isa, Quan Zou, Leyi Wei

**Affiliations:** School of Opto-electronic and Communication Engineering,Xiamen University of Technology,Xiamen, Fujian, China; School of Electrical and Electronic Engineering, Engineering Campus, Universiti Sains Malaysia, 14300 Nibong Tebal, Pulau Pinang, Malaysia; Yangtze Delta Region Institute (Quzhou), University of Electronic Science and Technology of China, Quzhou, Zhejiang, China; Institute of Fundamental and Frontier Sciences, University of Electronic Science and Technology of China, Chengdu, Sichuan, China; Faculty of Applied Sciences, Macao Polytechnic University, Macao 999078, China

## Abstract

Accurate prediction of effector proteins secreted by Gram-negative bacteria is important for elucidating bacterial pathogenic mechanisms and developing precise anti-infective strategies. Although existing methods have benefited from the strong sequence feature extraction capacity of pretrained protein language models, reliance on linear sequence information alone often fails to fully capture the three-dimensional conformational signals required for virulence functions. Meanwhile, conventional structure-based methods are limited by the scarcity of experimentally resolved protein structures. To address these challenges, We propose GeoEPred, a multimodal deep learning framework designed for the synergistic modeling of protein sequence and structure to identify Gram-negative bacterial effector proteins. Specifically, the model integrates sequence-contextual embeddings from a pretrained protein language model with three-dimensional structural representations predicted by ESMFold. A feature projection network refines fine-grained sequence signals associated with effector functions, while geometric vector perceptrons characterize inter-residue orientations, distances, and local spatial topology to capture potential structural conformational motifs. To further enable effective cross-modal fusion, we design a cross-modal alignment and feature-tokenized self-attention module. This module enhances consistency between the sequence-semantic and structural-geometric spaces through contrastive learning and models associations between linear functional motifs and spatial conformational patterns at a fine-grained token level. Extensive evaluations on multiple benchmark datasets show that GeoEPred achieves better predictive performance than existing leading models in T3SE, T4SE, and T6SE prediction tasks, while maintaining stable performance in remote homolog recognition scenarios. Moreover, the modular and extensible architecture of GeoEPred demonstrates strong generalization ability and substantial application potential for genome-scale effector protein discovery.

**Author summary:** Secreted effector proteins are central virulence factors used by many Gram-negative bacterial pathogens to execute infection strategies. Their functions are governed not only by secretion signals and short linear motifs in the amino acid sequence, but also by three-dimensional folds, local domains, and surface geometric patterns. However, current predictors mainly exploit sequence-contextual features, limiting their ability to model the correspondence between linear sequence signals and spatial conformational motifs, and thereby constraining accuracy and interpretability. Here, we present GeoEPred, a multimodal deep learning framework for secreted effector protein identification. GeoEPred couples sequence-semantic embeddings from a pretrained protein language model with structural representations learned by geometric vector perceptrons. A cross-modal alignment and interaction module uses contrastive learning to improve functional consistency between sequence and structure modalities, while feature-token attention captures fine-grained links between key linear and conformational motifs. Across benchmark datasets covering multiple effector types, GeoEPred outperforms existing state-of-the-art methods and provides interpretable evidence from sequence fragments, structural regions, and cross-modal associations, supporting functional annotation, pathogenic mechanism analysis, and experimental validation.

## Introduction

Gram-negative bacteria deploy specialized secretion systems to deliver virulence effector proteins that establish infection by modulating host signaling and evading immune responses [1–4]. Systematic discovery and functional profiling of these proteins are therefore essential for elucidating the molecular basis of pathogenesis and developing targeted antibacterial interventions. Although effectors are increasingly identified through experimental screening approaches, such as secretomic analyses [5, 6], these methods are often labor-intensive and time-consuming, and their outcomes can be influenced by variation in protein expression and secretion levels. To address these limitations, a range of computational tools has been developed, providing more efficient and cost-effective alternatives for effector protein identification.

Early computational tools for effector prediction largely relied on conventional feature engineering. For example, Bastion3 [7] and Bastion6 [8] integrated evolutionary conservation represented by position-specific scoring matrices (PSSMs) with structural constraint information, enabling effective prediction of type III and type VI effectors, respectively. CNN-T4SE combined PSSMs, protein secondary structure, solvent accessibility, and one-hot encoding to improve T4SE identification [9]. However, further optimization of such models is limited by the scarcity of annotated samples and by the substantial heterogeneity of effectors in sequence composition, secretion signals, and functional motifs, which constrains their ability to learn robust features from limited data [10].

In recent years, protein language models (PLMs) have provided new opportunities for effector protein prediction. By learning contextual representations enriched with evolutionary and functional information from large-scale protein sequences, PLMs can improve model generalization in low-data settings [11, 12]. Building on this advantage, DeepSecE [13] introduced a secretion-specific Transformer module to capture secretion-related patterns from PLM-derived sequence embeddings, whereas MoCETSE [14] combined a hybrid convolutional expert module with a Transformer incorporating relative positional encoding to further extract key discriminative features from PLM representations. Although these approaches have improved effector prediction, their inputs remain restricted to one-dimensional sequence information and do not explicitly exploit functional cues encoded in protein three-dimensional structures. Given that protein function is often jointly governed by tertiary conformation and spatial residue–residue interactions, sequence-based representations alone may be insufficient to fully characterize functionally diverse bacterial effectors [15–17].

A major reason for this limitation is the scarcity of experimentally resolved effector structures, which has hindered the systematic integration of structural features into prediction models. AlphaFold has transformed protein structure prediction [18], enabling large-scale access to high-quality structural models, and predicted structures have shown practical value in studies of intrinsically disordered proteins. However, the high computational cost of AlphaFold2 limits its application to genome-scale analyses. To address this issue, Lin et al. developed ESMFold [11], a structure prediction model that substantially improves inference efficiency while maintaining high prediction accuracy, accelerating structure prediction by up to approximately 60-fold. The efficiency of ESMFold makes large-scale protein structure analysis more feasible and provides a practical route for incorporating three-dimensional structural information into bacterial effector prediction [19, 20].

In this study, we propose GeoEPred, an end-to-end framework for effector protein prediction, designed to overcome the limitations of existing methods that rely heavily on sequence-derived features and insufficiently exploit protein spatial conformational information. GeoEPred first employs the pretrained ESM-2 model to extract protein embeddings, which are subsequently refined by a feature projection network to learn discriminative representations associated with virulence-related functions. To incorporate structure-level functional information, GeoEPred uses a geometric vector perceptron (GVP) module to extract three-dimensional conformational features from protein structures predicted by ESMFold. Unlike conventional graph neural networks that are restricted to scalar feature representations, GVP jointly models scalar and vector information, thereby capturing residue-level spatial relationships and local geometric orientations with higher fidelity. Furthermore, GeoEPred introduces a contrastive learning module to enhance cross-modal consistency between sequence semantic representations and structural geometric representations. It also employs a feature-tokenized multi-head attention mechanism to enable fine-grained interaction modeling, explicitly learning the latent associations between linear sequence signals and three-dimensional conformational motifs. GeoEPred not only outperforms existing mainstream binary classification models, but also achieves greater precision than widely used multiclass classifiers. The proposed method is also applicable to practical scenarios involving multiple types of effector proteins, enabling the simultaneous identification of diverse effectors without relying on explicit parameterization of physicochemical properties. Overall, our model provides a new perspective on the application of three-dimensional structural information to effector prediction and may help deepen our understanding of pathogenic mechanisms associated with bacterial secretion systems.

## Materials and methods

### Dataset construction

The datasets employed for model training and testing in this study were primarily sourced from the training and testing sets of established protein effector prediction models and related databases.

For T1SE and T2SE, we derived corresponding subsets from the TSE-ARF dataset [21]. Because these sequences had undergone stringent data curation and biological validation, they provided a reliable basis for building high-quality classification models. For T3SE and T6SE, sequence data were collected by integrating several resources, including Bastion3 [7], DeepT3 [22], SecReT6 [23], Bastion6 [8], DeepT3 4 [24], and TSE-ARF [21]. After merging these databases, we obtained a large pool of candidate effector protein sequences. The T4SE dataset was compiled from multiple prediction tools and databases, namely DeepSecE [13], OPT4e [25], T4SEfinder [26], iT4SE-EP [27], Bastion4 [28], T4SE-XGB [29], TSE-ARF [21], T4SEpp [30], and DeepT3 4 [24]. To reduce sequence redundancy and avoid model overfitting, redundant sequences were removed using CD-HIT [31] with a sequence identity cutoff of 60%. Dataset S1 summarizes the data sources and the sample sizes of each effector subtype after CD-HIT filtering. Based on the processed sequences, we constructed a multitask dataset, and the distribution of different effector categories is shown in S1 FigA. Specifically, the training set contains 1,575 non-effector proteins, 130 T1SEs, 68 T2SEs, 521 T3SEs, 675 T4SEs, and 272 T6SEs, whereas the test set includes 150 non-effector proteins, 20 T1SEs, 10 T2SEs, 30 T3SEs, 30 T4SEs, and 20 T6SEs. The proteins in this dataset span a broad length range, from fewer than 100 amino acids to more than 1000 amino acids, with most sequences concentrated between 100 and 700 amino acids (S1 FigB).

During the benchmarking stage, we selected appropriate benchmark datasets for different categories of secreted proteins in order to comprehensively assess the performance of GeoEPred in comparison with existing mainstream binary and multiclass prediction methods. For Type I secreted proteins, no publicly available independent test set is currently available. Therefore, following the strategy adopted in DeepSecE [13], we constructed a new benchmark dataset (Dataset S3) by extracting 20 T1SE samples and 150 non-T1SE samples from the independent test dataset S2. For Type III, Type IV, and Type VI secreted proteins, we directly employed the publicly accessible independent test datasets released by Bastion3, CNN-T4SE, and Bastion6, which were denoted as S4, S5, and S6, respectively. These benchmark datasets had been carefully compiled by the original authors using primary literature, public databases, and experimental evidence, they cover a wide range of bacterial species and research scenarios. Detailed information on all datasets used in this study is summarized in S1 Table.

### The architecture of the model

We propose GeoEPred, a novel approach that incorporates structural features to enhance effector protein prediction (Fig. 1A). Protein sequences are input into the pretrained protein language models ESMFold and ESM2, producing predicted structural PDB files and 2,560-dimensional sequence embeddings, respectively. The sequence embeddings are projected to 256 dimensions through a feature projection network. Concurrently, based on the predicted backbone structures, proteins are represented as residue-level neighbor graphs, in which each node corresponds to an amino acid residue and its spatial position is defined by the C*α* atom coordinates. Edges connect each residue to its nearest neighbors in three-dimensional space, with scalar and geometric vector features assigned to nodes and edges. A Geometric Vector Perceptron (GVP) is embedded in the graph neural network to jointly aggregate scalar and vector features from neighboring nodes and edges during message passing and feedforward updates, thereby capturing geometric relationships, local orientations, and residue interaction patterns. After aggregating these features into tokens, multimodal information is integrated via two modules: a contrastive learning module and a feature-token attention module. Contrastive learning aligns sequence semantic and structural geometric information in a unified representation space, enabling the model to capture their latent biological relationships. The feature-token mechanism partitions features into semantically distinct tokens and employs attention to model inter-token dependencies, allowing fine-grained interactions between sequence signals and structural conformations.

**Fig 1.**
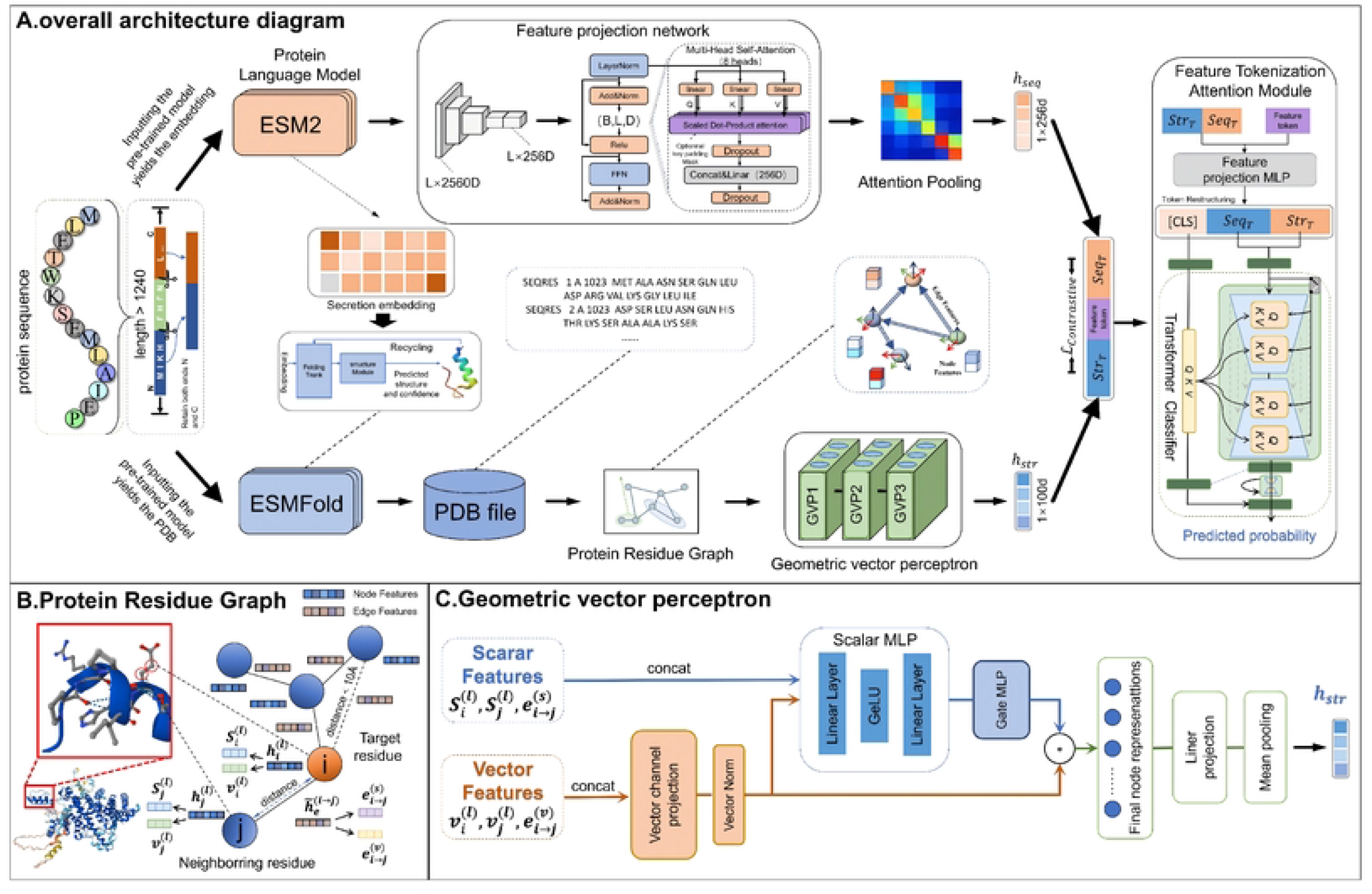
The overall workflow of GeoEPred. A: Effector protein sequences are initially encoded using a pre-trained protein language model to extract contextual evolutionary embeddings, which are subsequently refined through a feature projection network to generate sequence-level semantic representations. Concurrently, the input sequences are submitted to ESMFold to predict their three-dimensional conformations. Based on the predicted structures, protein residue graphs are constructed and processed with GVP-based message passing to learn geometry-aware structural representations. The resulting sequence semantics and structural features are then organized into distinct modality-specific feature tokens for downstream cross-modal representation learnin. B: Construction of Protein Residue Graph. C: Geometric Vector Perceptron.

### Protein language model

Previous studies have demonstrated that embeddings generated by protein language models contain rich information about protein sequence patterns and properties [32, 33]. This capability has promoted their widespread use in tasks involving protein sequence prediction and functional annotation. To determine the optimal protein language model (PLM) for sequence semantics, we evaluated several state-of-the-art architectures (S3 Table). We employ ESM-2 (t36) [11] with 650 million parameters to generate initial sequence semantic representations. During downstream training, the ESM-2 weights remain frozen to preserve the general biological patterns acquired from large-scale pre-training while reducing computational overhead. The parameter sources for each PLM are listed in S4 Table.

### Feature projection network

The raw ESM-2 embeddings *H*_esm_ ∈ ℝ^*L*×2560^ represent high-dimensional sequence features. We constructed a feature projection network to project these embeddings into a 256-dimensional latent space and model inter-residue dependencies:

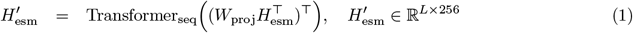

where *W*_proj_ ∈ ℝ^256×2560^ is the projection matrix for dimensionality reduction, and Transformer_seq_ captures long-range contextual patterns discriminative for effector identification.

Since the functional activity of effector proteins often relies on localized regions, such as N-terminal signal peptides or conserved motifs, global average pooling may dilute these critical site-specific signals. To address this, we develop a residue-adaptive attention pooling module that learns task-specific residue importance and selectively aggregates secretion- and virulence-related sequence representations. Specifically, let 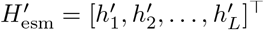 denote the set of refined semantic vectors for a protein sequence of length *L*, where each 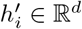 is the *d*-dimensional column vector representing the *i*-th residue. The importance score *a*_*i*_ for the *i*-th residue is defined as:

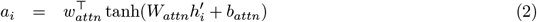

where *W*_*attn*_ ∈ ℝ^*k*×*d*^ and *b*_*attn*_ ∈ ℝ^*k*^ denote the weight matrix and bias vector that project the semantics into an attention space, *w*_*attn*_ ∈ ℝ^*k*^ is the weight vector that produces the raw importance scores (*d* = 256 and *k* = 64). The final global sequence semantic representation *h*_seq_ is obtained via a weighted sum of the residue features:

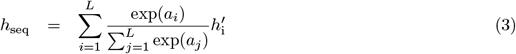

This mechanism prioritizes key functional segments while maintaining a comprehensive global context, providing a precise semantic input for subsequent multi-modal fusion.

### Protein Residue Graph Construction

To encode the three-dimensional structural information of proteins, each predicted protein structure is represented as a residue-level spatial graph *G* = (*V, E*) (Fig. 1B). Each node corresponds to an amino acid residue, with its spatial position defined by the *C*_*α*_ atom coordinate, and its initial representation consists of scalar and vector features describing local residue geometry. Scalar features include normalized backbone distances, the *N* − *C*_*α*_ − *C* bond angle and its sine, and amino acid type index. Vector features capture local residue orientation, including the unit vector from *C*_*α*_ to *C*, the normal vector of the backbone plane, and the corresponding orthogonal direction. The initial node representation 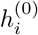 consists of scalar features 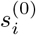 and vector features 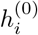, serving as input to the subsequent GVP-based structural encoder.

Edges are constructed based on spatial proximity between residues. An edge is added if two residues are within 10 Å *C*_*α*_ − *C*_*α*_ distance, and sequentially adjacent residues are always connected in both directions to preserve backbone continuity. Each directed edge *e*_*j*→*i*_ contains both scalar features 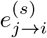 and vector features 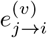, with scalar and vector features defined as:

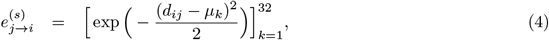

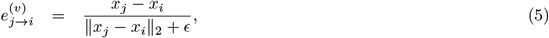

where *d*_*ij*_ is the Euclidean distance between residues *i* and *j, µ*_*k*_ is the *k*-th radial basis function center evenly spaced between 0 and 20Å, and *ϵ* is a small constant for numerical stability.

The resulting residue-level graph *G* includes node scalar features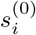, node vector features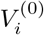, edge indices, and edge scalar and vector features.

### Geometric Vector Perceptron

To further encode the three-dimensional structural information of the residue graph, we implemented a Geometric Vector Perceptron (GVP)-based structural encoder [34]. By embedding GVPs into the graph message-passing process, the model generates messages from both node and edge geometric features, simultaneously updating scalar and vector node representations. This design enables the model to capture directional geometric information that is difficult to represent using scalar-only graph neural networks.

As illustrated in Fig. 1C, each residue node at layer *l* is represented as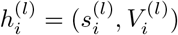, where 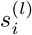 and 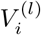 denote the scalar and vector features, respectively:

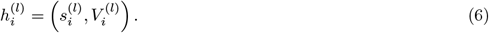

Each edge *j*→ *i* has embedded features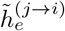, which also contain scalar and vector components. In each GVP-based graph convolution layer, messages are generated from the source node, target node, and edge features. Specifically, these features are integrated into a joint representation and passed into a GVP-based message function *g*^(*l*)^(·):

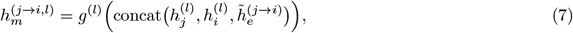

where 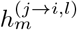 contains both scalar and vector messages. Within *g*^(*l*)^(·), vector features are transformed and coupled with scalar features through vector norms and scalar-dependent gating, enabling the generated message to encode both invariant and directional geometric information. Therefore, the information propagated along each edge depends not only on scalar residue representations, but also on geometric vector features such as edge directions, distance encodings, and local residue coordinate frames.

Incoming messages are aggregated by mean pooling and added to the current node representation through a residual connection:

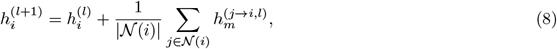

where 𝒩(*i*) denotes the set of incoming neighbors of node *i*. This mean aggregation reduces the influence of node-degree variation and enables stable integration of local neighborhood information.

After GVP-based message passing, the updated node representations encode geometric information through the joint propagation of scalar and vector channels. We then apply mean pooling over the final scalar node representations to obtain the protein-level structural embedding:

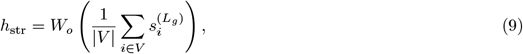

where *W*_*o*_ denotes the output linear projection, *V* is the set of residue nodes, and 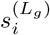 is the final scalar representation of residue *i* after *L*_*g*_ GVP-based graph convolution layers. Through the joint scalar–vector updates in GVP-based graph convolution layers, directional geometric information encoded by the vector channels is incorporated into the final node representations. Consequently, the structural embedding *h*_str_ captures both residue-level topological relationships and three-dimensional geometric context, and is subsequently used as the structural representation for multimodal fusion and downstream prediction tasks.

### Cross-modal feature tokenization

After obtaining the protein structural embedding *h*_str_ from the GVP module, the model integrates it with the sequence embedding *h*_seq_ produced by the sequence encoder. To enhance the interaction between sequence and structure modalities, the two global embeddings are first normalized and projected into a shared *d*-dimensional latent space to construct modality-level feature tokens. In addition, independent cross-modal projection layers are used to generate a coupled interaction token through element-wise multiplication:

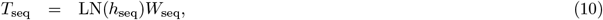

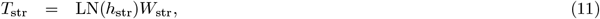

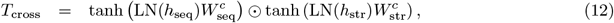

where *W*_seq_ and *W*_str_ are learnable projection matrices for the sequence and structure modalities, respectively. 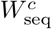 and 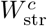 denote independent projection matrices for constructing the cross-modal ⊙ interaction token, and denotes element-wise multiplication. The resulting *T*_seq_, *T*_str_, and *T*_cross_ represent the sequence feature token, structural feature token, and cross-modal interaction token, respectively.

A learnable classification token *T*_cls_ is then concatenated with the three feature tokens. Positional embeddings *E*_pos_ are added to the token sequence, which is subsequently fed into a multi-layer Transformer encoder. The Transformer self-attention mechanism models dependencies among the sequence-specific, structure-specific, and cross-modal interaction representations:

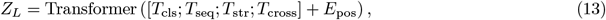

where *Z*_*L*_ denotes the final token representations after *L* Transformer encoder layers. The output corresponding to the classification token, denoted as *z*_cls_, aggregates global cross-modal context and is passed to a classification head to generate the final prediction.

During training, the model is optimized using a joint objective that combines the classification loss and the cross-modal contrastive loss. The classification loss ℒ_cls_ is implemented as Focal Loss with label smoothing, while the contrastive loss ℒ_con_ adopts the NT-Xent objective to encourage consistency between sequence and structure representations in the projection space:

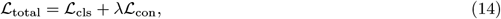

where *λ* controls the relative contribution of the contrastive learning objective.

By applying Transformer-based self-attention to the sequence, structure, and coupled interaction tokens, the model can emphasize discriminative modality-specific and cross-modal representations, thereby forming a unified protein representation for downstream effector classification and secretion-system type prediction.

## Results

### Evaluation of GeoEPred Performance in Effector Protein Prediction

GeoEPred was evaluated through five-fold cross-validation using training and validation subsets derived from dataset S1. Furthermore, the model’s generalization capability was verified on an independent dataset S2. Using five-fold cross-validation, the model achieved an average accuracy of 88.43%, demonstrating robust performance across folds. On the independent test set, GeoEPred also performed well, achieving an accuracy of 93.85%. To more intuitively reflect the overall performance, we plotted a confusion matrix, shown in Fig. 2B and Fig. 2D, which displays the number of correctly and incorrectly predicted samples for each class. The confusion matrix indicates that the model correctly predicted a large number of samples across the five effector protein classes.

**Fig 2.**
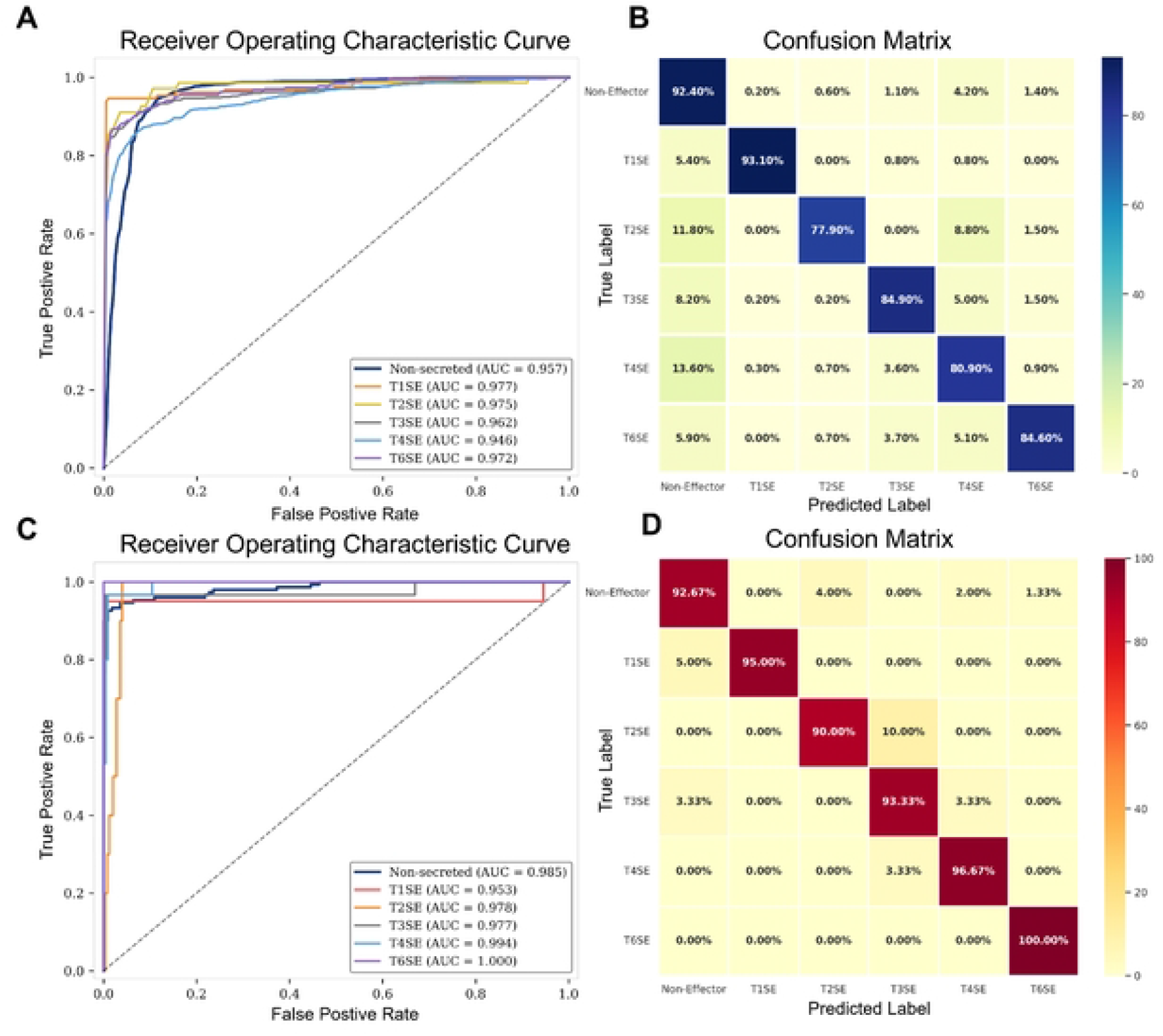
Assessing the predictive efficacy of GeoEPred in identifying bacterial secretion system effectors. A-D: Model performance assessed via five-fold cross-validation (A, B) and independent testing (C, D). ROC curves illustrate the classification performance for each effector type. Sensitivity values for each class are indicated along the diagonals of the corresponding confusion matrices.

To systematically validate the model’s discriminative ability across different effector protein classes, ROC curves were generated(Fig. 2A,Fig. 2C). The AUC values for all categories exceeded 0.9, demonstrating robust inter-class discrimination ability in multi-class scenarios. This quantitative evidence confirms the model’s superior classification performance and operational reliability. Notably, despite the smaller sample size for the T2SE category, its corresponding ROC curve still exhibits high discriminative power.

### Ablation study of characteristic sources and fusion strategies

To assess the contribution of each modality and evaluate the synergistic effect of the multimodal fusion module, we performed a comprehensive ablation study. All methodological variants were subjected to a uniform training-validation pipeline with standardized hyperparameters and metrics to ensure the integrity of cross-comparisons.

#### Multimodal feature ablation

Specifically, we decoupled the network architecture across modalities and conducted a detailed examination of the individual contributions of purely structural features, purely sequence features, refined sequence semantics, and the final multimodal fusion features to the performance of effector protein prediction. Table 1 provides a detailed quantitative ablation analysis of each feature configuration on the validation set. Experimental results show that multimodal fusion of sequence semantics with structural features yields the optimal predictive performance (ACC = 93.85%, MCC = 90.61%), significantly surpassing all single-modality baselines. A deeper comparative analysis reveals that relying solely on purely structural features yields limited discriminative capability (ACC 50.76%); in contrast, although using sequence semantics alone achieves substantial performance gains (ACC = 88.08%, SN = 87.06%), it still exhibits obvious blind spots when handling more complex biological patterns. By introducing a structural features, the multimodal collaborative framework achieves an MCC gain of 8.27% on top of sequence semantics baseline, effectively overcoming the intrinsic limitations of pure sequence models. This result provides strong empirical support for a high representational complementarity between evolutionary sequence semantics and spatial geometric features. The fusion mechanism, by leveraging the joint advantages of both modalities, expands the model’s decision boundary and markedly enhances predictive robustness.

**Table 1.** Performance evaluation of multimodal feature ablation.

| Feature Type | ACC (%) | AUC | SN (%) | MCC | SP (%) |
| --- | --- | --- | --- | --- | --- |
| Structure only | 50.76 | 82.30 | 50.67 | 32.27 | 89.05 |
| Sequence only | 70.38 | 92.19 | 72.11 | 60.10 | 94.03 |
| Sequence Semantics | 88.08 | **98.48** | 87.06 | 82.34 | 97.57 |
| GeoEPred | **93.85** | 98.44 | **94.61** | **90.61** | **98.72** |

To validate the above quantitative findings, Fig. 3 employs Uniform Manifold Approximation and Projection (UMAP) [35] to visualize the latent feature space of validation samples under different feature configurations. When sequence or structural features are used in isolation, sample embeddings exhibit substantial dispersion, with severe feature overlap between positive and negative classes, directly leading to high misclassification rates. Conversely, introducing ESM embedding alone markedly improves cluster homogeneity, but the centers of all clusters remain relatively close, causing samples near the decision boundary to be easily confused. Although after refining embedded representation to condense information, a clearer separation emerges between effector and non-effector proteins, inter-effector classification remains suboptimal. In stark contrast, the joint fusion of sequence semantics and structural features yields the most discriminative topology: positive and negative samples form dense clusters in the high-dimensional embedding space with well-defined, mutually separated boundaries. This visual evidence robustly confirms that cross-modal complementarity expands the decision boundary; from a manifold-learning perspective, it explains the substantial performance gains.

**Fig 3.**
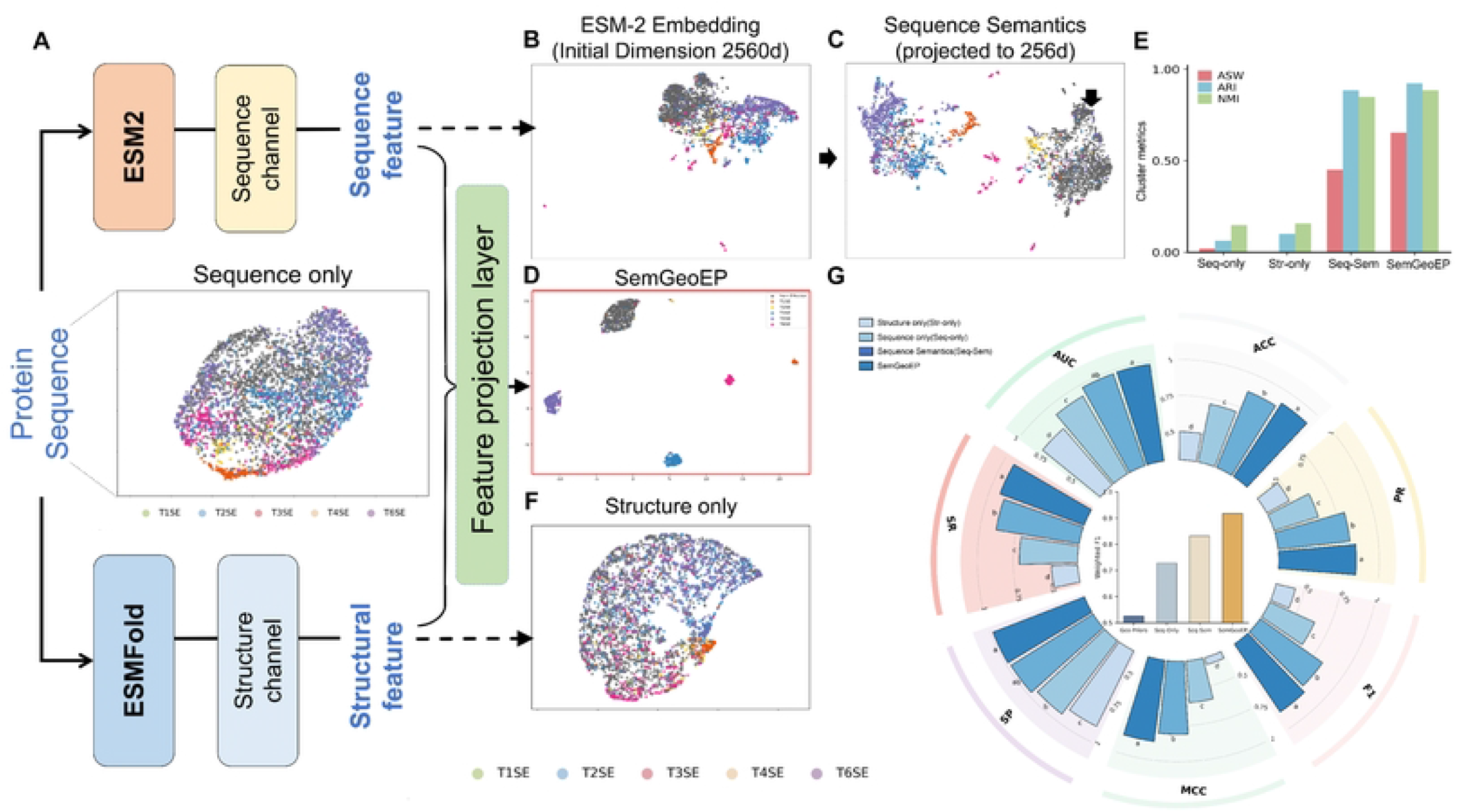
Ablation Analysis and Latent Space Visualization of GeoEPred. A: Multi-modal Input Streams: Schematic representation of the sequence features (ESM-2 embeddings and refined sequence semantics) and the structural features in isolation. B: UMAP visualizations of effector and non-effector protein clusters based on ESM-2 embedding vectors (2560 dimensions). C: Refined Sequence Semantics: UMAP visualization of protein clusters after sequence-channel projection and contextual encoding (256 dimensions). D: Multimodal UMAP of Effectors: UMAP visualization showing the precise clustering of effectors and non-effectors after integrating both sequence semantics and structural features. E: I compare clustering performance with bar plots against the evaluation metrics NMI, ARI, and ASW. F: Structural features: UMAP visualization of effector and non-effector clusters derived from the GVP. G: A ring diagram compares ablation metrics, with the center representing the weighted F1.

Three clustering performance metrics—Adjusted Rand Index(ARI), Normalized Mutual Information(NMI) and Average Silhouette Width(ASW)—further indicate that modality fusion substantially enhances the clustering capability of the output representations(Fig. 3E). Compared with a single modality, GeoEPred shows improvements in ARI and NMI, suggesting a stronger ability to discriminate among different effectors.

#### Precise classification of proteins with distant homology

In biochemistry, protein function is primarily determined by its three-dimensional structure rather than solely by its amino acid sequence [36]. Notably, some proteins can maintain highly similar spatial structures despite low sequence identity, thereby exhibiting similar functional properties. Compared with conventional sequence alignment methods, structural alignment is capable of capturing functional relationships across greater evolutionary distances [37]. In general, protein pairs with sequence identity below 0.3 and TM-score above 0.5 are considered to exhibit significant structural similarity and are often regarded as remote homologs [38].

To further validate the necessity of incorporating structural information, we performed a focused ablation study under the extreme “long-range homology” scenario. Specifically, we curated a dedicated benchmark of 107 proteins (S4 Fig), each satisfying the remote-homology criterion with respect to the training set. As shown in Fig. 4, the structures of samples in the training set and the independent test set have a certain degree of overlap in 3D space.

**Fig 4.**
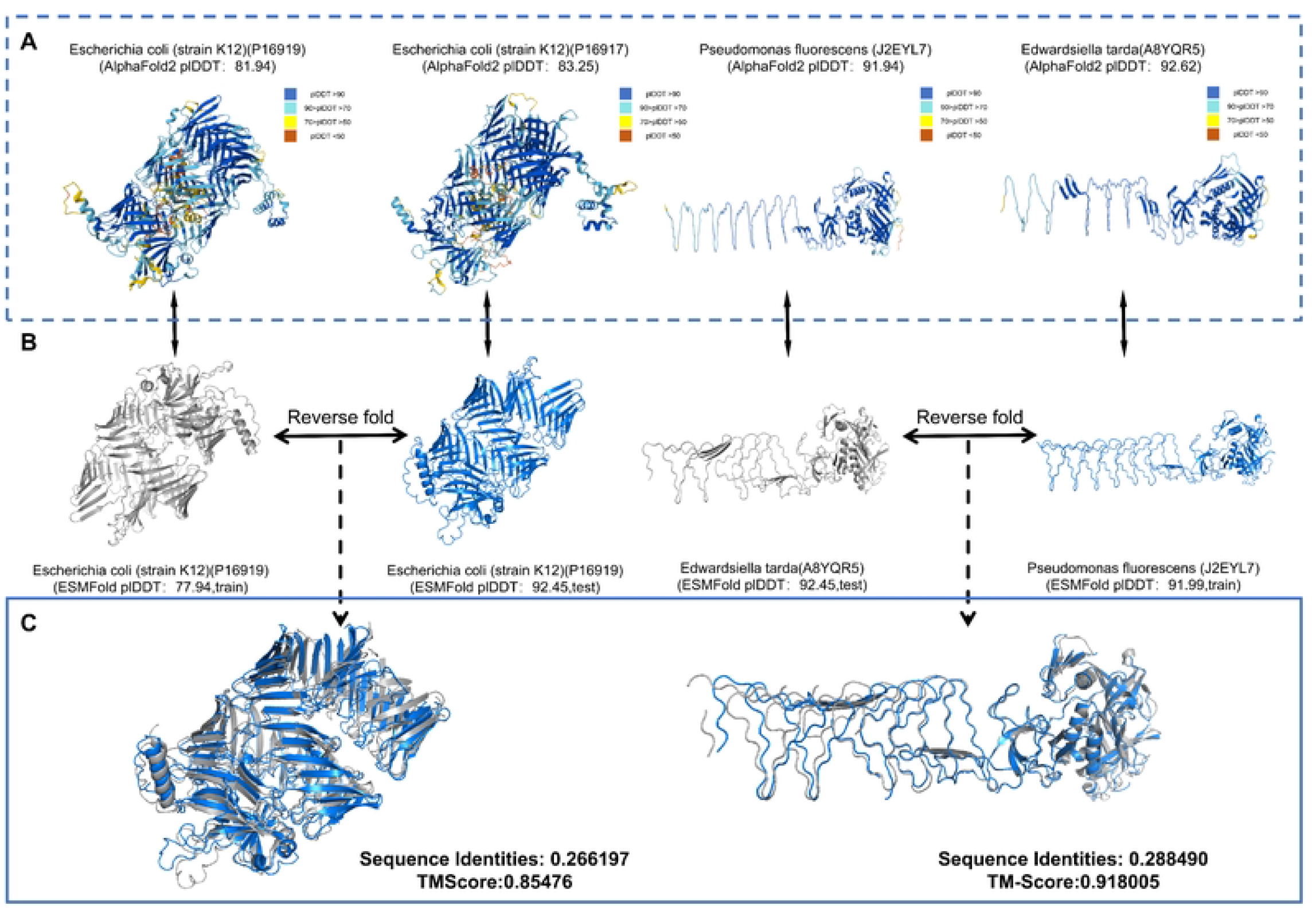
The degree of spatial overlap between the 3D structures of distant homologs. A: Three-dimensional protein structures predicted by AlphaFold2, colored according to pLDDT scores (higher scores indicate higher confidence). B: Three-dimensional protein structures predicted by ESMFold. C: The middle panel displays the structural alignment of two proteins from each group. “Train” indicates samples from the training set, while “Test” indicates samples from the test set. Sequence identity represents the percentage of aligned amino acid positions that are identical between two protein sequences.TM-score quantifies the structural similarity between two protein structures, with higher scores indicating greater similarity.

On this benchmark, we compared GeoEPred with two ablated variants: one relying on sequence semantics refined by the feature projection network alone, and the other using only raw sequence features.

The raw-sequence variant performs poorly, reaching an accuracy of just 65.42%, and misclassifies 23 of the 38 T3SE samples into other categories. Substituting raw features with sequence semantics produces a marked improvement, raising the number of correctly classified samples to 88. This gain shows that contextual representations from pre-trained protein language models, coupled with the feature projection network that distills fine-grained functional signals, markedly reduce the model’s reliance on superficial sequence similarity. GeoEPred further lifts the accuracy to 90.65% (S6 Table), delivering the best performance on this benchmark. The result confirms that integrating refined sequence semantics with geometric structural features is the principal driver of performance gains in this challenging scenario.

### Ablation research on fusion mechanism

Following the confirmed effectiveness of multimodal feature fusion, we further examine the extent to which distinct feature integration strategies influence model performance. An ablation study is carried out that benchmarks three representative fusion paradigms under a unified experimental condition: Feature Tokenization—the core mechanism adopted in GeoEPred—together with Cross-Attention Fusion and Feature Concatenation. To guarantee that the comparison is fair and reproducible, all strategies are trained and validated under identical procedures, share the same hyperparameter configuration, and are evaluated with a common set of assessment metrics.

As reported in Table 2, Feature Tokenization consistently outperforms the two competing fusion strategies across all five evaluation metrics. Specifically, it surpassing both Cross-Attention Fusion and Feature Concatenation by a clear margin. Compared with Cross-Attention Fusion, which yields the lowest overall performance (ACC = 89.62%, MCC = 84.35%), Feature Tokenization delivers absolute gains of 4.23% in ACC and 6.26% in MCC, indicating substantially stronger discriminative capability. Feature Concatenation performs slightly better than Cross-Attention Fusion (ACC = 90.00%, AUC = 98.32%), yet still falls short of Feature Tokenization by 3.85% in ACC and 5.38% in MCC. Notably, the improvement in MCC—a metric that is particularly sensitive to class imbalance—suggests that Feature Tokenization not only enhances overall predictive accuracy but also produces more balanced predictions across positive and negative samples. These results collectively confirm that token-level integration of multimodal features provides a more effective fusion mechanism than either attention-based interaction or naive concatenation within the GeoEPred framework.

**Table 2.** Performance evaluation metrics for different fusion strategies.

| Strategy | ACC (%) | AUC | SN (%) | MCC | SP (%) |
| --- | --- | --- | --- | --- | --- |
| Cross-Attention Fusion | 89.62 | 97.63 | 86.67 | 84.35 | 97.95 |
| Concatenation | 90.00 | 98.32 | 91.89 | 85.23 | 98.01 |
| Feature Tokenization | **93.85** | **98.44** | **94.61** | **90.61** | **98.72** |

### Interpretable investigation into multimodal feature contributions

Building upon the validated efficacy of multimodal feature fusion, we performed SHAP (SHapley Additive exPlanations) analysis grounded in cooperative game theory. This method quantifies feature importance via a combined global and local interpretability metric, illustrating how multimodal features jointly contribute to prediction outcomes.

The class-specific feature importance rankings are illustrated in Fig. 5A, in which the top 20 features are listed in descending order of their mean SHAP values. Notably, ESM-derived sequence semantics (e.g., esm f85, esm f144) dominate the highest-ranked positions, indicating that evolutionary sequence information plays a decisive role in driving the model’s predictions. These features encode the conserved biochemical and functional semantics embedded in homologous sequences, thereby providing strong discriminative signals for effector identification. Meanwhile, structural features also contribute substantially, with at least one structural feature entering the top 20 across all categories—an effect that is particularly pronounced in T3SE and T6SE (e.g., struct f10, struct f18). This observation highlights the complementary role of the structural feature, which functions analogously to a spatial constraint filter: it injects spatial structural information that sequence features alone struggle to characterize, thereby reducing false-positive predictions caused by sequence-level ambiguity.

**Fig 5.**
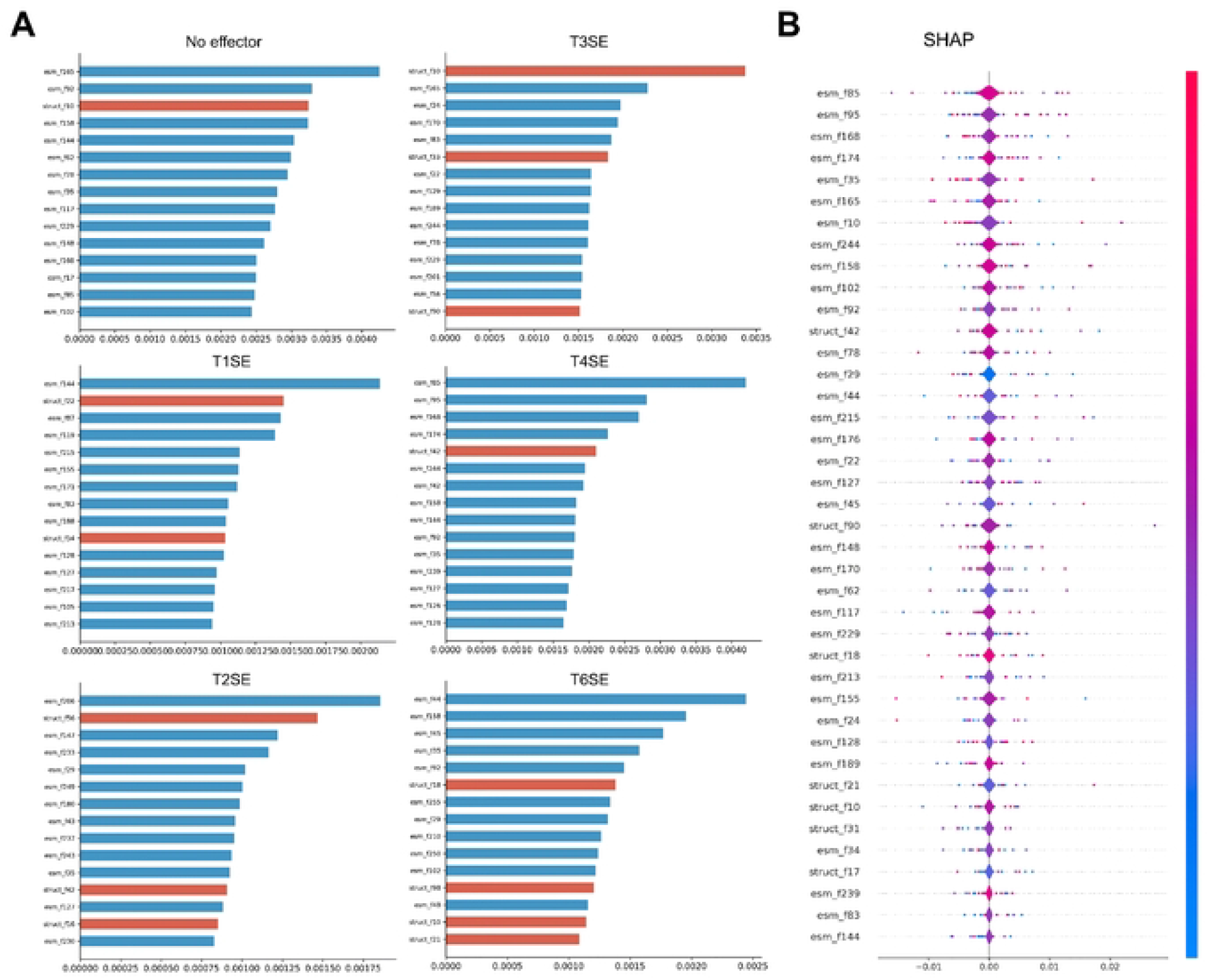
Feature Importance Rankings by Category and SHAP Swarm Plots. A: Ranking of feature importance for individual categories. B: Ranking of overall features and SHAP analysis

The SHAP summary plot (Fig. 5B) further elucidates the relationship between feature values and their impact on model output. Leading sequence features (e.g., esm_f85, esm_f95) exhibit a broad distribution of influence, with their values spanning a wide range of SHAP values from strongly positive to strongly negative. This bidirectional influence reflects their role in capturing complex, context-dependent evolutionary semantics, allowing them to support classification decisions under specific sequence patterns. In contrast, high-contribution structural features display a more concentrated pattern with a clearer directional tendency. For instance, several structural features (e.g., struct_f21, struct_f42) tend to yield positive SHAP values when their values are high, suggesting that such spatial constraints serve as key determinants for distinguishing certain effectors.

In summary, GeoEPred effectively exploits the division-of-labor advantages between distinct feature modalities: sequence semantics provide the core discriminative signal for effector identification, whereas the structural features offers finer-grained feature characterization for challenging samples. This multimodal fusion strategy successfully integrates the complementary information from both sources.

### Comparison with baseline methods

To comprehensively evaluate the architectural efficacy of GeoEPred, we performed a systematic benchmark against 11 baseline models under a uniform input configuration. These baselines encompass seven conventional machine learning algorithms (SVM, Random Forest, Gradient Boosting, KNN, Naive Bayes, Decision Tree, and LDA) and four representative deep learning architectures (MLP, BiLSTM-Attention, GRU, and a Vanilla Transformer).

As shown in the Table 3, GeoEPred achieved the best or most competitive results across the main evaluation metrics. Specifically, it reached accuracies of 93.85% (ACC), 90.61% (MCC), 94.61% (Recall), and 90.93% (F1-Score), outperforming all baselines; its precision reached 88.45%, slightly below Random Forest’s 89.09%, but remained highly competitive overall. These results indicate that GeoEPred not only exhibits stronger overall classification performance but also excels particularly in identifying positive samples.

**Table 3.** Comparison of the proposed GeoEPred method with baseline methods.

| Method | ACC (%) | MCC (%) | Rec (%) | F1 (%) | Pr (%) |
| --- | --- | --- | --- | --- | --- |
| *Traditional Machine Learning* |  |  |  |  |  |
| SVM | 90.77 | 86.08 | 91.83 | 87.76 | 85.83 |
| Random Forest | 89.62 | 83.12 | 76.67 | 80.56 | **89.09** |
| Gradient Boosting | 86.92 | 79.33 | 75.06 | 74.90 | 77.17 |
| KNN | 83.08 | 76.99 | 88.17 | 80.40 | 79.27 |
| Naive Bayes | 81.54 | 74.20 | 82.94 | 75.69 | 74.06 |
| Decision Tree | 69.23 | 55.38 | 65.28 | 61.09 | 58.83 |
| LDA | 61.92 | 55.36 | 79.44 | 64.06 | 59.66 |
| *Deep Learning Methods* |  |  |  |  |  |
| BiLSTM-Attention | 90.00 | 85.36 | 92.33 | 86.36 | 83.25 |
| MLP | 90.00 | 85.24 | 90.06 | 84.99 | 82.38 |
| GRU | 90.00 | 85.45 | 92.33 | 86.03 | 82.77 |
| Vanilla Transformer | 88.05 | 84.15 | 92.72 | 85.23 | 81.79 |
| *Our method* |  |  |  |  |  |
| GeoEPred | **93.85** | **90.61** | **94.61** | **90.93** | 88.45 |

Further comparisons reveal that among traditional machine-learning methods, SVM’s overall performance is the closest to ours, yet its ACC, MCC, Recall, and F1-Score are approximately 3.08, 4.53, 2.78, and 3.17 percentage points lower than GeoEPred, respectively. Among the deep-learning baselines, BiLSTM Attention, and GRU perform comparatively well, but they still lag behind ours across the composite metrics. These results demonstrate that GeoEPred’s architecture is superior to conventional models in capturing complex discriminative patterns.

### Comparison with state-of-the-art models

To rigorously evaluate GeoEPred’s efficacy in identifying Gram-negative secretion effectors (T1SE–T4SE and T6SE), we benchmarked it against three multi-class frameworks—MoCETSE [14], DeepSecE [13], BastionX [39], and six specialized binary classifiers using datasets S3–S6. Under uniform testing conditions, predictions for baseline methods were retrieved from their official web servers. Specifically, binary models were assessed on their respective target datasets (S3 for T1SEstacker [40]; S4 for Bastion3 [7] and T3SEpp [41]; S5 for Bastion4 [28], T4SEfinder [26], and T4SEpp [30]; S6 for Bastion6 [8]), while multi-class models were evaluated across the entire benchmark suite. Performance was quantified using ACC, REC, PR, F1-score, and MCC, with tool access details provided in S2 Table.

From the overall results, GeoEPred exhibits superior predictive capabilities compared to current state-of-the-art approaches, a trend most evident in T3SE, T4SE, and T6SE prediction tasks. In the T1SE prediction task, GeoEPred demonstrated superior performance compared to T1SEstacker (S2 Fig), which relies on amino acid composition features without RTX C-terminal motifs. The accuracy improved from 92.9% to 98.2%, the F1 score from 72.7% to 94.7%, and the MCC from 69.1% to 94.2% (S4 FigA). For T3SE, GeoEPred maintains a high level of predictive performance, with ACC 98.1% and F1 91.8%, while PR reaches 90.3%, clearly outperforming MoCETSE, DeepSecE, and T3SEpp on this metric. In the comparison of multi-class classification models, although MoCETSE successfully predicted 29 T3SE effectors, it produced a higher number of false positives compared to GeoEPred (as shown in Fig. 6A). This indicates that GeoEPred can sustain strong overall classification performance while also reducing false positives.In the T4SE task, GeoEPred’s advantages are more pronounced. The model achieves the best results with ACC 96.2%, F1 92.3%, and MCC 90.5%, all superior to MoCETSE (ACC 95.4%, F1 90.9%, MCC 88.7%), DeepSecE (ACC 95.4%, F1 90.6%, MCC 88.0%) and Bastion4 (ACC 91.5%, F1 84.1%, MCC 79.3%), as illustrated in Fig. 6B. Although multiple models can effectively identify T4SE effectors, GeoEPred performs better in terms of false positive rate (Fig. 6D). DeepSecE and MoCETSE achieved satisfactory classification results on the T6SE prediction task, while GeoEPred performed exceptionally well, with ACC, F1, and MCC all reaching 1.000 (Fig. 6C), surpassing DeepSecE (96.8%, 96.7%, 93.6%) and MoCETSE (98.6%, 92.7%, 92%). (For detailed performance information of each model, please refer to S5 Table.)

**Fig 6.**
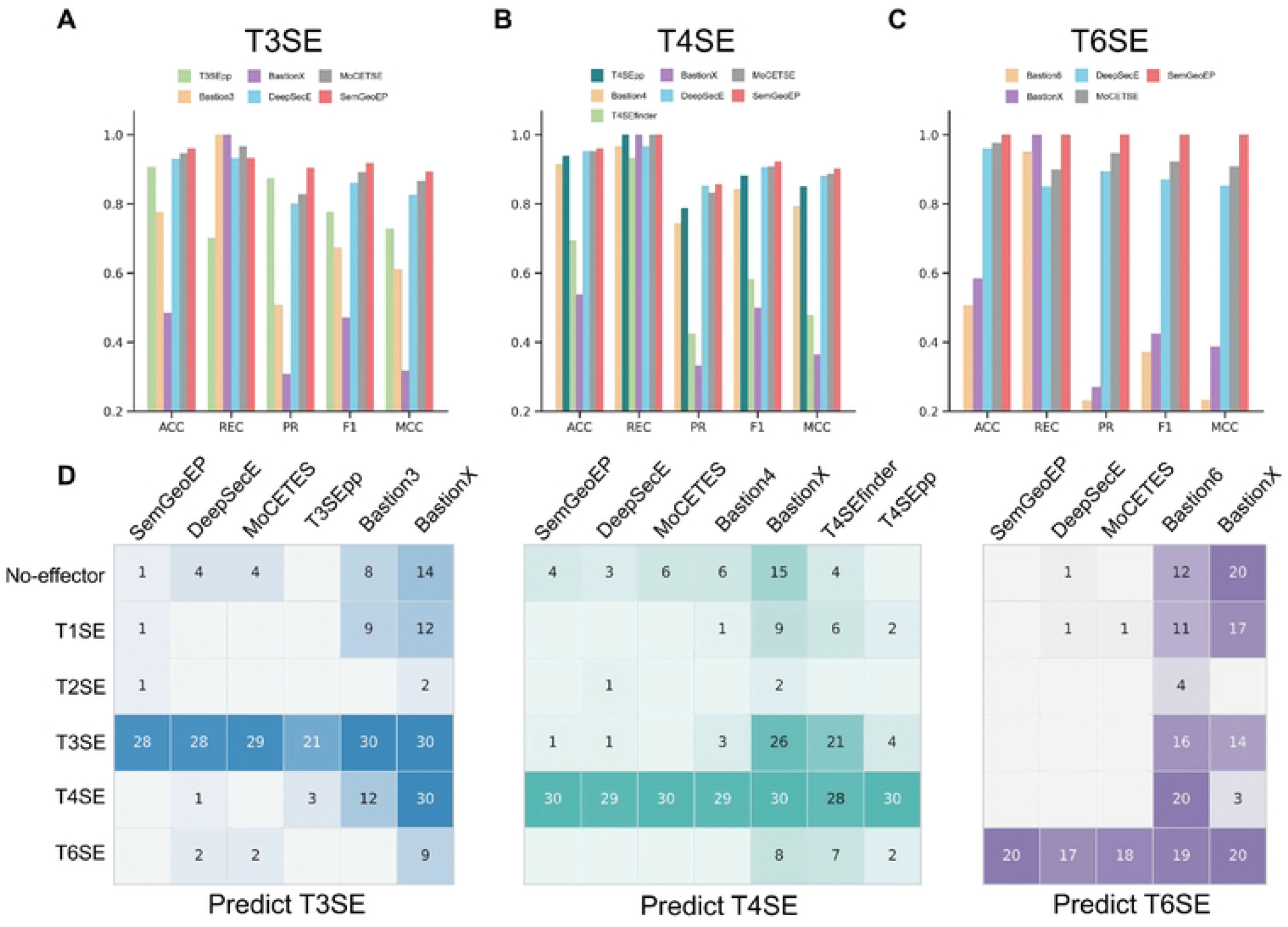
Performance evaluation of GeoEPred in comparison with mainstream models. A-C: Performance comparison of GeoEPred with mainstream prediction models on T3SE (A), T4SE (B), and T6SE (C) prediction tasks. D: Comparison of true positive and false positive predictions from different models on the independent test set (Dataset S2), which contains 150 non-effectors, 20 T1SEs, 10 T2SEs, 30 T3SEs, 30 T4SEs, and 20 T6SEs. The heatmaps illustrate the distribution of predicted samples across various effector protein types.

While some binary methods may achieve higher sensitivity on specific tasks, they can entail higher cross-class false-positive risk, particularly in multiclass independent test sets. Compared with several binary approaches, GeoEPred yields fewer false positives for other types, suggesting that a single binary predictor may be less suited than a multiclass model for distinguishing a given type from others(Fig. 6B). From a practical standpoint, because a single bacterial genome typically encodes multiple types, multiclass models with well-balanced performance across categories offer greater utility. In this regard, GeoEPred demonstrates superior stability and balance in multiclass recognition, making it more suitable for reliable predictions in complex genomic backgrounds.

Finally, the confusion-matrix analyses indicate that GeoEPred achieves low false-positive rates in both binary and multiclass settings. For example, in T4SE, T4SEpp, a SOTA binary model, can correctly identify all T4SEs and non-T4SE proteins but misclassifies other types. By contrast, because we incorporated other types as negatives during training, GeoEPred achieves a PR of 1.000 for T4SE, a property that may translate into more reliable predictions in more complex genomic contexts.

### case studies

Genome-wide prediction of secreted effector proteins in Gram-negative bacteria is a crucial approach for investigating their pathogenicity, distribution patterns, and evolutionary dynamics [42]. Compared with labor-intensive and time-consuming experimental methods, bioinformatics approaches enable more efficient screening of potential effector proteins, providing a practical solution for large-scale studies of their distribution, evolutionary trajectories and pathogenic potential [10]. To assess the practical applicability of GeoEPred, we developed a computational pipeline that integrates MacSyFinder’s specialized secretion system models [43] with the GeoEPred framework. Based on this integration, we developed a computational pipeline that predicts secretion systems and their associated secreted proteins directly from bacterial genome data, thereby enabling subsequent prediction of secretion substrates.

We applied GeoEPred to perform genome-wide screening of secreted proteins from 5,571 protein-coding sequences of *Pseudomonas aeruginosa PAO1* (NCBI accession: NC 002516.2), successfully identifying gene clusters corresponding to Type III and Type VI secretion systems. To further evaluate the capability of GeoEPred as a genome-wide prediction model, we specifically analyzed the predicted Type VI secreted effectors in *Pseudomonas aeruginosa PAO1*. GeoEPred correctly identified the majority of experimentally validated effectors (28 out of 30), achieving a recall of 93.3%. Meanwhile, the high-confidence candidate proteins identified by GeoEPred, including T1SE, T2SE, and T6SE (S3 Fig), show strong genomic proximity to the secretion system loci detected by MacSyFinder (Fig. 7), suggesting that these candidate effectors may be functionally and genomically associated with their corresponding secretion machinery. This observation is consistent with previous reports showing that some experimentally validated effector proteins are frequently located near secretion system gene clusters or pathogenicity islands [44].

**Fig 7.**
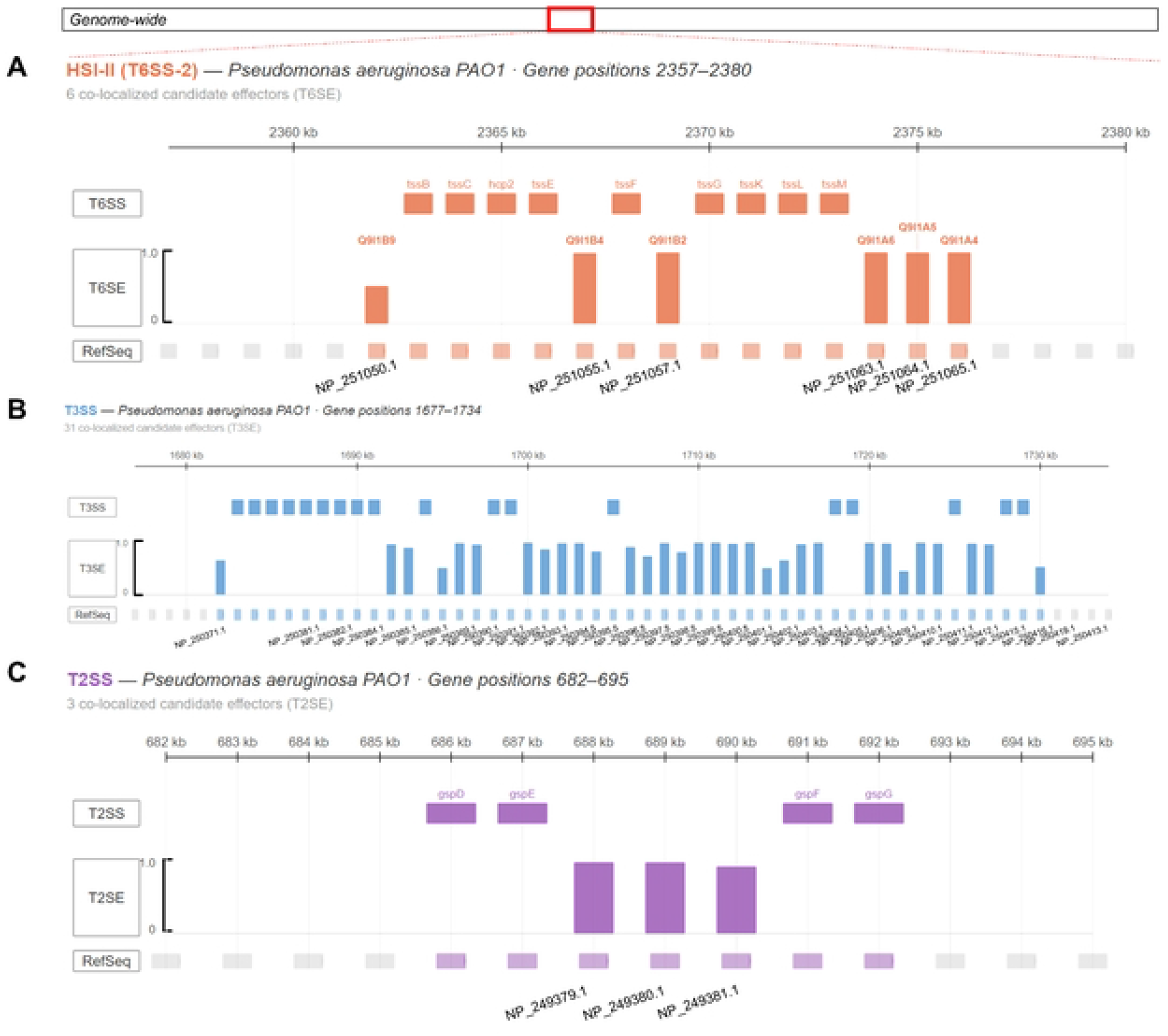
Genome-wide predictions indicate that effector proteins are distributed near the secretory system.

In addition, we further evaluated the model on two other Gram-negative bacterial genomes: *Salmonella enterica subsp. enterica serovar Typhimurium str. LT2* (NC 003197.2) and *Legionella pneumophila subsp. pneumophila str. Philadelphia 1* (NC 002942.5). The predicted results were compared against experimentally validated data. Genome reliably identified a large proportion of experimentally confirmed T3SE and T4SE, achieving recall rates of 94.78% (36 out of 38) and 91.2% (280 out of 307), respectively. These results show that GeoEPred can perform well across different biological contexts. It also has strong robustness and reliability in Large-scale genome-wide annotation tasks.

### Biological interpretability

To interpret the biological signals learned by GeoEPred, we visualized the prediction results using sequence significance maps and PyMOL [45], allowing us to examine the model’s decision patterns from both residue-level motif information and three-dimensional structural conformations. At the residue level, the feature projection network refined contextual information to identify key amino acid segments associated with effector secretion and delivery, particularly characteristic signal regions at the N- and C-termini.

Complementarily, at the structural level, the GVP module extracted spatial residue proximity and local geometric orientation from ESMFold-predicted structures, enabling the model to capture three-dimensional functional cues that are difficult to infer from linear sequences alone.

In the analysis of the *Salmonella typhimurium* T3SS SlrP, GeoEPred first localized the N-terminal region (Fig. 8D), whose first 50 residues encode the T3SE translocation signal [46, 47]. Beyond this linear delivery signal, the structural branch showed a strong preference for C-terminal residues 751–765 (Fig. 8A), which correspond to the key interface through which NEL-family effectors recruit host E2 ubiquitin-conjugating enzymes [48, 49]. This indicates that structural features provide complementary spatial functional information beyond sequence-derived signals.

**Fig 8.**
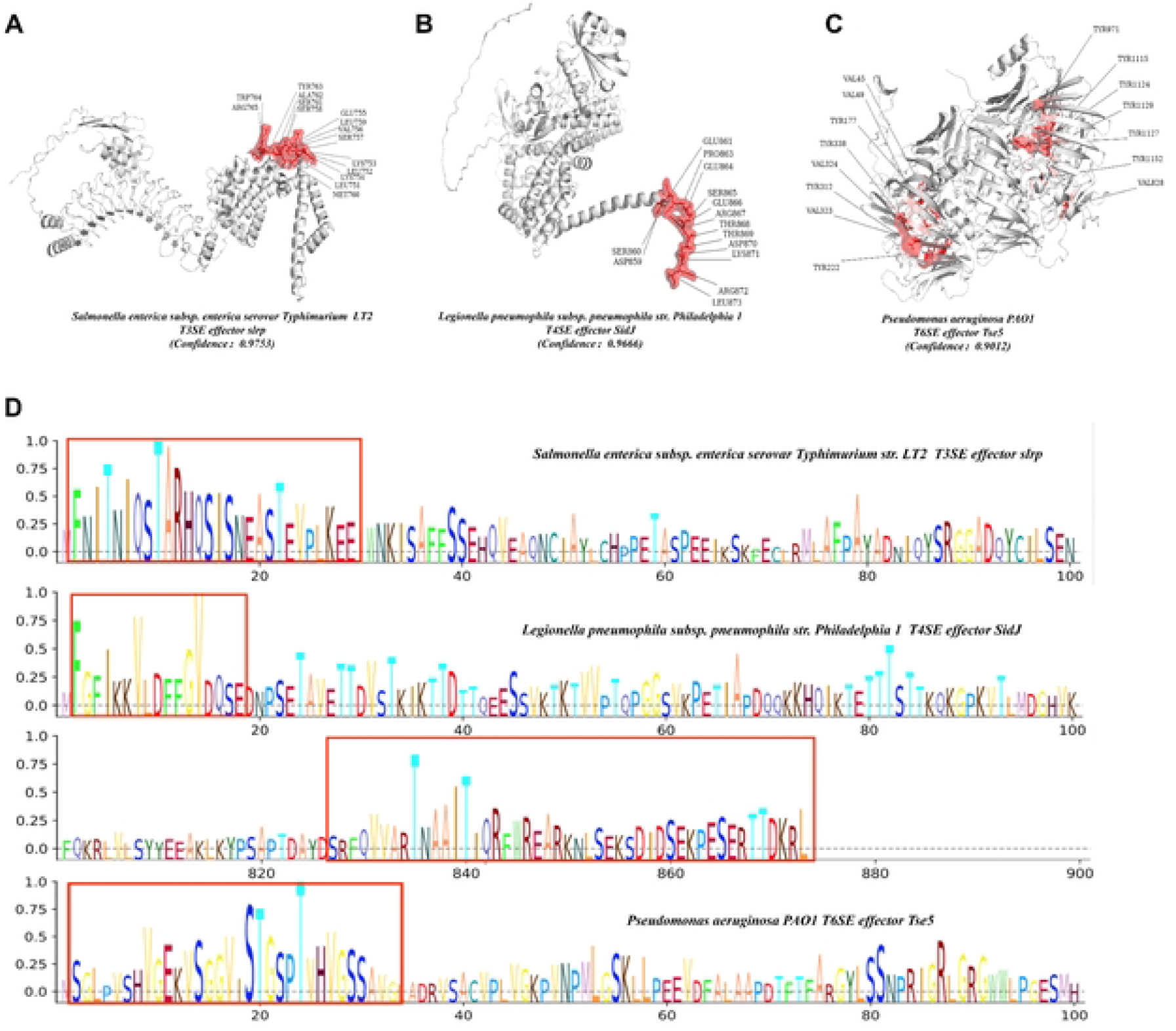
Performance comparison between GeoEPred and state-of-the-art predictors for bacterial secretion effectors. A-C: Visualization of the weights of E3 generalized enzyme Slrp (A), calmodulin-dependent gamma-glutamyltransferase SidJ (B), and toxin protein Tse5 (C). D: Visualization of the N-terminal and C-terminal sequences of the three enzymes, showing the amino acid preferences in their secretion patterns.

For the *Legionella pneumophila* T4SS effector SidJ, both sequence and structural visualizations converged on the C-terminal region (Fig. 8D, residues 859–873). This region is biologically meaningful because the C-terminal extension of SidJ is essential for binding host calmodulin (CaM), a prerequisite for activating its glutamylase activity [50, 51]. Thus, in SidJ, the two modalities provide convergent evidence by highlighting the same functional regulatory region from the perspectives of linear sequence patterns and three-dimensional conformation.

Regarding the *Pseudomonas aeruginosa PAO1* T6SS effector Tse5, the sequence branch mainly focused on the N-terminal region, residues 1–35 (Fig. 8D). This segment mediates interactions with Hcp and the iron-binding module of the PQS system and is a key determinant for Tse5 recruitment and secretion through the T6SS machinery [52]. In contrast, the structural branch was highly sensitive to the C-terminal region of the Rhs shell, residues 828–1152 (Fig. 8C), accurately identifying the toxin-encapsulation interface [53]. This finding agrees with high-resolution cryo-EM studies showing that the Rhs shell forms a cocoon-like protective structure that encloses the pore-forming toxin fragment Tse5-CT, which mediates target-cell membrane depolarization [54]. Overall, these results show that structural features do not simply duplicate sequence signals, but supplement linear sequence representations with critical spatial functional information, allowing GeoEPred to capture secretion and delivery cues, host-interaction interfaces, and toxicity-associated conformational features.

## Discussion

The application of deep learning to effector protein prediction in Gram-negative bacteria provides computational advantages that are difficult to achieve using conventional experimental approaches, offering new opportunities to investigate bacterial virulence mechanisms and support antibacterial drug design [10, 42, 55]. In this study, we present GeoEPred, an end-to-end model that captures both linear sequence semantics and 3D structural conformations for effector prediction. Unlike existing methods that rely primarily on sequence information, GeoEPred uses a geometric vector perceptron module to model three-dimensional conformational features, thereby addressing the limitations of sequence-only representations in capturing spatial functional information and improving predictive performance. Benchmark results show that GeoEPred achieves perfect performance on the T6SE task and outperforms representative existing methods on both T3SE and T4SE prediction(Fig. 6).

Further analyses demonstrate that the cross-modal contrastive learning module and feature-tokenized attention mechanism promote complementary integration between sequence semantics and structural geometry, substantially enhancing the overall discriminative capacity of the model (Table 2). ROC curves and confusion matrices indicate that GeoEPred maintains balanced performance across effector classes, supporting its robustness for multiclass classification (Fig. 4). Notably, GeoEPred also performs well in challenging scenarios involving remote homologs, further highlighting the importance of incorporating structural features for robust effector prediction (S6 Table).

Beyond accurate prediction, GeoEPred provides biologically interpretable evidence from both sequence and structural perspectives that is consistent with experimental observations. For example, the model highlights key signal regions at the N- and C-termini, as well as structural segments associated with virulence-related functions (Fig. 8). These results suggest that GeoEPred not only achieves strong predictive performance but also provides interpretable insights into effector protein function.

At the genome scale, secreted protein prediction typically requires the integration of homology-based inference and multiple feature-driven machine learning strategies [42]. In contrast, GeoEPred offers an efficient end-to-end framework for directly predicting secretion systems and their associated substrate proteins. We further applied GeoEPred to the genomes of three representative Gram-negative pathogens and evaluated its predictions against experimentally validated proteins. The results show that GeoEPred accurately identifies validated effectors and uncovers candidate secretion system-associated proteins from complex genomic backgrounds, providing useful clues for future effector discovery (Fig. 7, S5 Table).

Despite its strong performance and application potential, GeoEPred has several limitations. First, class imbalance among effector subtypes remains unresolved and could be further mitigated through targeted data augmentation or resampling strategies. Second, model stability may be improved by enhancing structural quality or incorporating additional sequence-derived features. Finally, extending GeoEPred to other secretion systems, such as T5SS and T7SS, as well as to Gram-positive bacteria, will help evaluate its applicability and generalizability across broader biological contexts.

## Supporting information

**S1 Appendix. Evaluation indicators**.

**S1 Fig. Visualization of protein sequence length distributions in the dataset**.

**S2 Fig. Performance comparison of GeoEPred and mainstream models in type I (T1SE) and type II (T2SE) secretion effector protein prediction tasks**. A-B: Model performance was evaluated on the benchmark set (Dataset S3) and a dataset containing 150 non-effect proteins and 10 T2SE types, using metrics such as accuracy (ACC), recall (REC), precision (PR), F1 score (F1), and Matthews correlation coefficient (MCC). D-E: Comparison of true positive and false positive predictions for T1SE and T2SE in different models. The heatmap shows the sample distribution predicted as T1SE, T2SE, and other effector protein types.

**S3 Fig. Genome-wide prediction visualization shows that GeoEPred not only successfully identifies T6SEs in Pseudomonas aeruginosa, but also predicts other candidate effectors in the genome (A–E)**.

**S4 Fig. The dataset contains samples with distant homology, including 2 T1SE, 42 T3SE, 34 T4SE, 9 T6SE, and 20 effector proteins**.

**S1 Table. Summary of the training, testing, and benchmarking datasets used in the comparative experiments**.

**S2 Table. S2 Table. Web server links of the comparative models used in this study**.

**S3 Table. Compares the performance of GeoEPred on independent test sets based on ESM-1b, ESM-2, and other protein language models**.

**S4 Table. GitHub repositories of protein language models evaluated for GeoEPred**.

**S5 Table. Performance comparison between GeoEPred and existing popular methods on benchmark datasets**.

**S6 Table. S6 Table. Ablation Comparison in Datasets Containing Remote Homologs**.

## Acknowledgments

This work was supported by the National Natural Science Foundation of China (No. 62372392), and the Graduate Scientific Innovation Project of Xiamen University of Technology (No. YKJCX2025082).

## Data availability

The data, code, and models from this study are publicly available on GitHub (https://github.com/shouzhensong/GeoEPred). This library enables researchers to access datasets, utilize the code, invoke the developed models, and make predictions.

## Notes

### Competing Interest Statement

The authors have declared that no competing interests exist.

